# Minor QTLs mining through the combination of GWAS and machine learning feature selection

**DOI:** 10.1101/702761

**Authors:** Wei Zhou, Emily S. Bellis, Jonathan Stubblefield, Jason Causey, Jake Qualls, Karl Walker, Xiuzhen Huang

**Affiliations:** Department of Computer Science and Molecular Bioscience Program, Arkansas State University, Jonesboro, Arkansas, 72401; Department of Biology, The Pennsylvania State University, University Park, Pennsylvania, 16802; Department of Computer Science and Mathematics, University of Arkansas at Pine Bluff, Pine Bluff, Arkansas 71601

**Keywords:** Machine learning, Minor QTLs, GWAS, Feature selection

## Abstract

**Introduction:** Minor QTLs mining has a very important role in genomic selection, pathway analysis and trait development in agricultural and biological research. Since most individual loci contribute little to complex trait variations, it remains a challenge for traditional statistical methods to identify minor QTLs with subtle phenotypic effects. Here we applied a new framework which combined the GWAS analysis and machine learning feature selection to explore new ways for the study of minor QTLs mining.

**Results:** We studied the soybean branching trait with the 2,137 accessions from soybean (*Glycine max*) diversity panel, which was sequenced by 50k SNP chips with 42,080 valid SNPs. First as a baseline study, we conducted the GWAS GAPIT analysis, and we found that only one SNP marker significantly associated with soybean branching was identified. We then combined the GWAS analysis and feature importance analysis with Random Forest score analysis and permutation analysis. Our analysis results showed that there are 36,077 features (SNPs) identified by Random Forest score analysis, and 2,098 features (SNPs) identified by permutation analysis. In total, there are 1,770 features (SNPs) confirmed by both of the Random Forest score analysis and the permutation analysis. Based on our analysis, 328 branching development related genes were identified. A further analysis on GO (gene ontology) term enrichment were applied on these 328 genes. And the gene location and gene expression of these identified genes were provided.

**Conclusions:** We find that the combined analysis with GWAS and machine learning feature selection shows significant identification power for minor QTLs mining. The presented research results on minor QTLs mining will help understand the biological activities that lie between genotype and phenotype in terms of causal networks of interacting genes. This study will potentially contribute to effective genomic selection in plant breeding and help broaden the way of molecular breeding in plants.

## Introduction

In molecular genetics research, a remaining challenge in quantitative trait studies is the efficient mapping of minor quantitative trait loci (QTLs) to identify causative genes and understand the genetic basis of variation in quantitative traits [1]. Because the subtle influence on the phenotype of minor QTLs is easily masked by epistasis [2] and gene-environment interactions [3], minor QTLs are more difficult to be detected and analyzed. Because of this, a large fraction of the genetic architecture of most complex traits is not well understood [4, 5, 6]. Currently, almost all of genes or QTLs that have been verified were major effect ones, and the minor effect QTLs were less investigated. Several different methods have been reported to identify minor QTLs, but many of these strategies have had poor success rates [7, 8, 9]. To improve the situation, some of these studies were based on expensive experimental data from large populations. For example, Baobao et al., demonstrated a method for mapping of minor effect QTLs in maize by using super high density genotyping and large recombinant inbred population [10].

QTL-mapping algorithm based on statistical machine learning methods better estimates of QTL effects, because it eliminates the optimistic bias in the predictive performance of other QTL methods. It produces narrower peaks than other methods and hence identifies QTLs with greater precision [17]. Two machine-learning algorithms (Random Forest and boosting) have been used to analyze discrete traits in a genome-wide prediction context. It was found out that Random Forest and boosting do not need an inheritance specification model and may account for non-additive effects without increasing the number of covariates in the model or the computing time [18]. This study shows some advantages in the use of machine learning methods to analyze discrete traits in genome-wide prediction. Random Forest was shown to outperform other methods in the field datasets, with better classification performance within and across datasets. Even when tested with the main QTLs for several traits in different chromosomes, Random Forest was able to identify them, but it failed to detect significant associations when the variance explained by the QTL is low [19].

Besides physical QTLs mapping, machine learning methods are also used on eQTL(Expression quantitative trait loci) Mapping. By using combinations of methods, an approach that relies on Random Forests and LASSO was developed and it achieved a much higher average precision at the cost of slightly lower average sensitivity [20]. It is observed that when combined Random Forest and other modeling techniques, it almost always performed better than their constituent methods [21, 22]. It is observed that Random Forests map eQTL are to be validated by independent data, when compared to competing multi-locus and legacy eQTL mapping methods [20].

Genome-wide association studies (GWAS) is considered to be a powerful approach for dissecting complex traits [23,24,25] and has been widely applied for the study of many plants, such as *Arabidopsis*, rice and maize [26, 27, 28, 29, 30, 31]. In soybean, the evaluation of several specific agronomic traits, including seed protein and oil concentration [32, 33], cyst nematode resistance [34, 35], and flowering time [36] were conducted through GWAS. Plant architecture related traits (PATs) are of great importance for soybean and many crops. Studies in past decades indicated that PATs are mainly affected by minor effect quantitative traits loci (QTLs), especially as reflected in the Nested Association Mapping (NAM) population [37, 38, 39].

From these previous studies, however, minor QTLs are hard to be detected mainly because their contribution is subtle. It is challenging for current statistical methods to detect them. For example, most of statistic methods are based on the variance analysis, such as ANOVA, and they usually need a larger population size to detect minor QTLs.

In this study, with soybean branching as the focused trait, we combined the GWAS analysis and machine learning feature selection, to explore the application of a new analysis framework in minor QTLs mining in plants. As a result, we identified 328 minor genes and 1770 effective SNP markers related to soybean branching development. Our analysis results with the new framework for minor QTLs mining would benefit the genomic selection, the pathway analysis and organism development research.

## Methods

### 1. Dataset

The original genotypic data is from soybase data bank: https://soybase.org/snps/. The SoySNP50K iSelect BeadChip has been used to genotype the USDA Soybean Germplasm Collection [46]. The complete data set for 20,087 G. max accessions genotyped with 42,509 SNPs is available. Soybean accessions and phenotypic data used in this study were obtained from the USDA Soybean Germplasm Collection (http://www.ars-grin.gov/npgs/). Branching phenotype data was extracted and used for analysis. Missing data and SNPs with minor allele frequencies below 0.1 were excluded, leaving 42,080 SNPs for GWAS.

### 2. Genome wide association study (GWAS)

Association analysis and estimation of each SNP effect was implemented in GAPIT software (version 2) [47]. The regression linear model (GLM), and the mixed linear model (MLM) methods were used as described by Tang et al. [45]. Default parameters of the SUPER model were used: sangwich.top = “MLM,” sangwich.bottom = “SUPER,” LD = 0.1. The significant P-value cut-off was set as p = 3.45e-07, equivalent to α level of 0.05 after Bonferroni correction. The efficient mixed-model association with corrections for kinship and population structure was applied. Three PCs generated from GAPIT were included as covariates. The SNPs with a minor allele frequency (MAF) higher than 0.01 were used to estimate the population structure and the kinship. Only SNPs with a MAF higher than 0.1 were used for association tests. The cutoff of significant association was a False Discovery Rate (FDR) adjusted P-value less than 0.1 using the Benjamini and Hochberg procedure to control for multiple testing. Significant SNPs were defined if showing a minus log10-transformed P ≥ 3. SNPs with a genetic distance less than 2 cM were considered to be in a LD extension block and belong to the same SNP cluster.

### 3. Data preprocessing

In machine learning feature selection analysis, all of nucleotides in genotype data was added the rs (Reference SNP cluster ID) information and transformed as rs + nucleotide (Sup_Table7). The whole dataset was divided into 11 subsets based on different P-value levels for a further analysis in machine learning models. The genotype data used in regression and feature importance analysis were encoded by OneHotEncoder after labelencoding.

### 4. Feature importance analysis

Feature importance analysis explains what features have the biggest impact on predictions in testing model. Permutation importance is a kind of global model-agnostic method and calculated after a model has been fitted. Compared to most other approaches, permutation importance is fast to calculate and widely used. Random forest is one of the most effective machine learning models for predictive analytics capable of performing both regression and classification tasks and able to capture non-linear interaction between the features and the target. It is very good at handling categorical features with fewer than hundreds of categories [49]. The character of permutation importance consists with the properties we would want a feature importance measure to have. In this research we applied the random regressor in permutation importance analysis and Random Forest score analysis for all of 2137 samples and 42080 features (SNPs).

### 5. Gene Ontology analysis

SNPs identified by feature importance analysis were searched in SoyBase data site (https://soybase.org/snps/) by rs number. And the flank sequence of corresponding SNP was used to BLAST in Glycine max Genome DB database (http://www.plantgdb.org/GmGDB/) for confirmation. The gene names which SNPs hit to the same location (including CDS, UTR and intron) were collected for GO (gene ontology) analysis. All the genes identified by BLAST were analyzed by GO term enrichment tool at SoyBase website (https://soybase.org/goslimgraphic_v2/dashboard.php). The GO enrichment information, related charts and gene location map were generated by GO term enrichment tool at SoyBase website.

## Results

### 1. Genome Wide Association Study (GWAS) for soybean branching

A genome-wide association study (GWAS) of soybean branching was conducted with 42,080 SNP markers in the GAPIT (Genome Association and Prediction Integrated Tool) software using a mixed-linear model (MLM). 3541 SNP markers with P-value less than 1.0 were identified. Among these 3541 markers, there are 18 markers with P-value less than 0.005, 32 markers with P-value less than 0.01 and 161 makers with P-value less than 0.05(Table 1. and Sup_Table1.). Associations between phenotypes and genetic markers are displayed as Manhattan plots (Fig. 1) and (Sup_Table1). P-values were displayed in negative log scale with base of 10 (–log10 (P)) against the physical map positions of genetic markers. We set a threshold of −log10 (0.1/42080) = 5.624 (42080 is the SNP marker numbers) to identify SNPs significantly associated with a trait. In total of 161 which P-value is less than 0.05, only SNP marker ss715607451 were significantly (−log10 (p) = 9.524328812) associated with soybean branching trait. Marker ss715632223 and ss715613636 with log10 (p) value at 4.634512015 and 4.554395797 respectively, are near to the threshold but not reach it (Fig. 1; Sup_Table1). In other words, by the GAPIT analysis, only one SNP marker significantly associated with soybean branching was identified. We also BLAST the 18 SNPs which P-value less than 0.005 in Soybase and five annotated genes are found (Table 1), but none of them is reported as branching related.

**Table 1.**
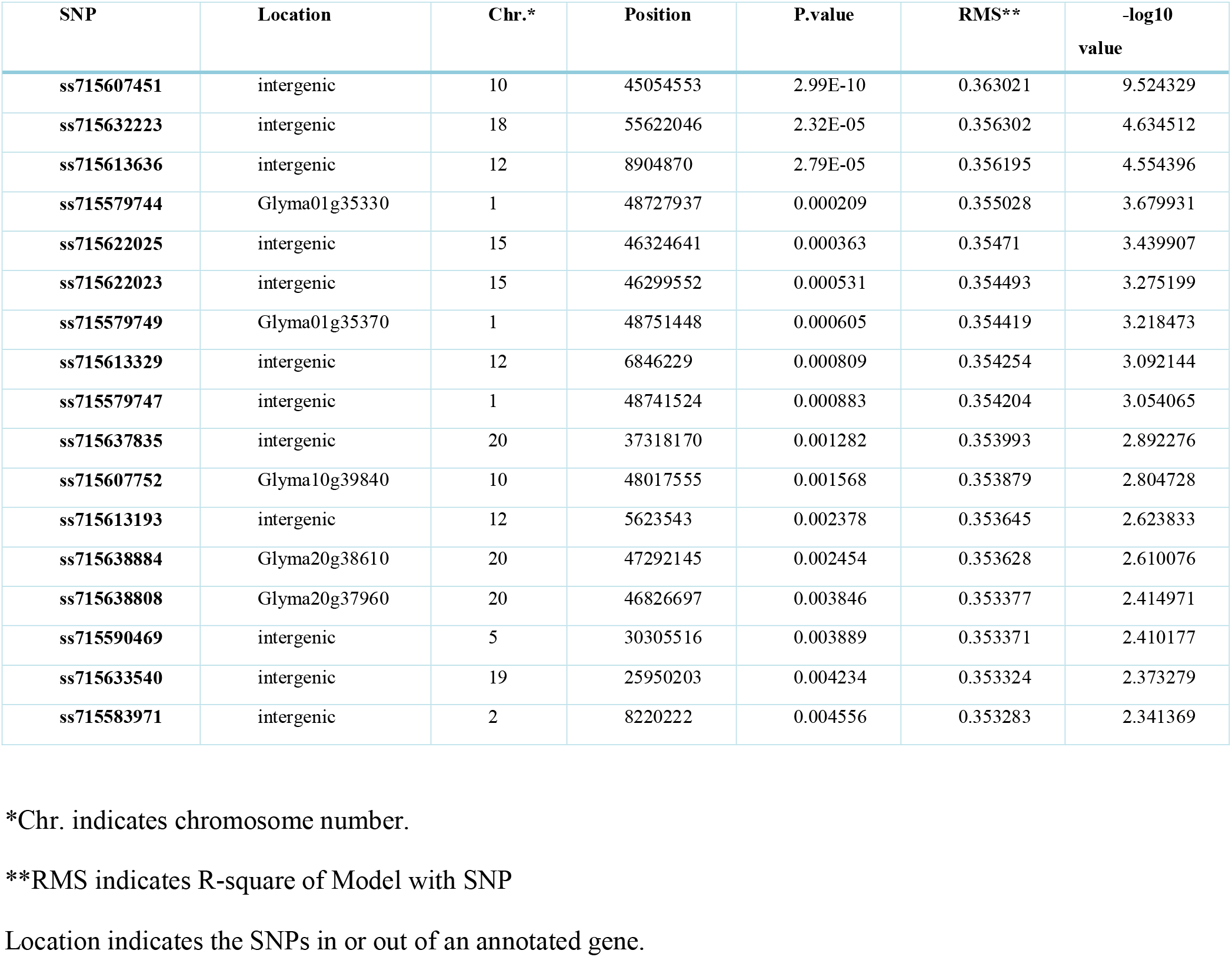
Summary of SNP markers with P-value less than 0.005 from GWAS analysis.

| SNP | Location | Chr.* | Position | P.value | RMS** | -log10<br>value |
| --- | --- | --- | --- | --- | --- | --- |
| ss715607451 | intergenic | 10 | 45054553 | 2.99E-10 | 0.363021 | 9.524329 |
| ss715632223 | intergenic | 18 | 55622046 | 2.32E-05 | 0.356302 | 4.634512 |
| ss715613636 | intergenic | 12 | 8904870 | 2.79E-05 | 0.356195 | 4.554396 |
| ss715579744 | Glyma01g35330 | 1 | 48727937 | 0.000209 | 0.355028 | 3.679931 |
| ss715622025 | intergenic | 15 | 46324641 | 0.000363 | 0.35471 | 3.439907 |
| ss715622023 | intergenic | 15 | 46299552 | 0.000531 | 0.354493 | 3.275199 |
| ss715579749 | Glyma01g35370 | 1 | 48751448 | 0.000605 | 0.354419 | 3.218473 |
| ss715613329 | intergenic | 12 | 6846229 | 0.000809 | 0.354254 | 3.092144 |
| ss715579747 | intergenic | 1 | 48741524 | 0.000883 | 0.354204 | 3.054065 |
| ss715637835 | intergenic | 20 | 37318170 | 0.001282 | 0.353993 | 2.892276 |
| ss715607752 | Glyma10g39840 | 10 | 48017555 | 0.001568 | 0.353879 | 2.804728 |
| ss715613193 | intergenic | 12 | 5623543 | 0.002378 | 0.353645 | 2.623833 |
| ss715638884 | Glyma20g38610 | 20 | 47292145 | 0.002454 | 0.353628 | 2.610076 |
| ss715638808 | Glyma20g37960 | 20 | 46826697 | 0.003846 | 0.353377 | 2.414971 |
| ss715590469 | intergenic | 5 | 30305516 | 0.003889 | 0.353371 | 2.410177 |
| ss715633540 | intergenic | 19 | 25950203 | 0.004234 | 0.353324 | 2.373279 |
| ss715583971 | intergenic | 2 | 8220222 | 0.004556 | 0.353283 | 2.341369 |

\*Chr. indicates chromosome number.
\*\*RMS indicates R-square of Model with SNP
Location indicates the SNPs in or out of an annotated gene.

**Fig. 1.**
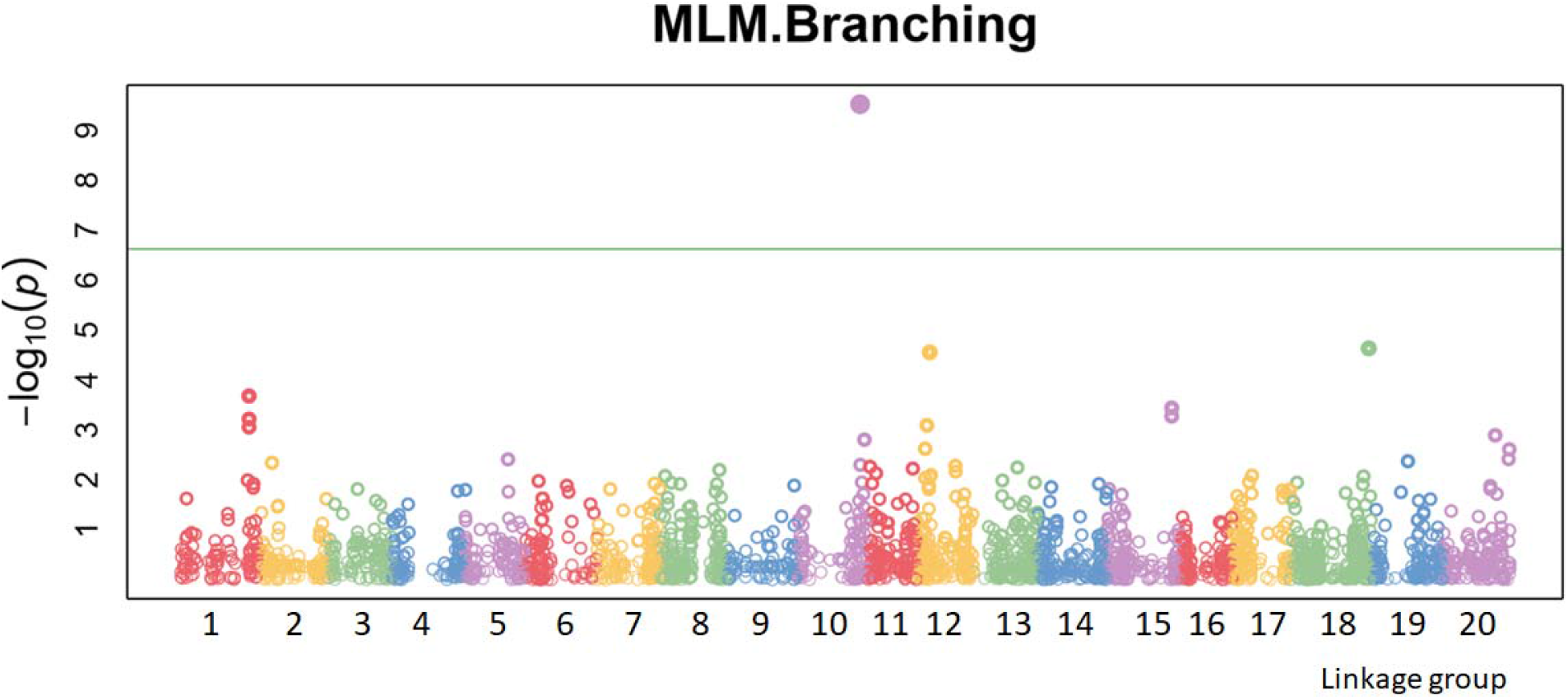
Manhattan plots of genome-wide association studies (GWAS) for soybean branching. Manhattan plots of genome-wide association studies (GWAS) for soybean branching measured with the mixed linear model (MLM). The X-axis is the genomic position of the SNPs in each linkage group, and the Y-axis is the negative log base 10 of the P-values. Each chromosome is colored differently. SNPs with stronger associations with the trait will have a larger Y-coordinate value. The general and highly significant trait-associated SNPs are distinguished by the green threshold lines. Genetic markers are positioned by their chromosomes and ordered by their base-pair positions. Genetic markers on adjacent chromosomes are displayed with different colors. The strength of the association signal is displayed in two ways. One indicator of strength is the height on the vertical axis for −log P-values; the greater the height, the stronger the association. The other indicator is the degree of filling in the dots; the greater the area filled within the dot, the stronger the association.

### 2. Feature importance analysis

Please refer to Fig. 3 for a summary chart of our feature importance analysis. In the following we give the details of our analysis results.

In general, feature importance analysis is based on the understanding how the features in the testing model contribute to the prediction model. Feature importance includes local model-agnostic feature importance and global model-agnostic feature importance. Since local measures focus on the contribution of features for a specific prediction, whereas global measures take all predictions into account. Here we applied permutation feature importance, a global model-agnostic approach, with the Random Forest algorithm as the core. After evaluating the performance of the models, we permuted the values of a feature of interest and re-evaluate the model performance. The average reduction in impurity across all trees in the forest due to each feature was computed.

Our results showed that there are 974 features in total with the weight values above zero. Among them, 971 features (SNPs) have weights bigger than 1E-06, 952 features (SNPs) in total have weights bigger than 1E-05 and 872 features (SNPs) have weights bigger than 0.0001(Sup_Table2.). Our results also showed that there are 1124 features in total with negative weight values. Among them, 1107 features (SNPs) have weights smaller than −1E-05, 939 features (SNPs) have weights smaller than 1E-04 (Sup_Table2.). There are 39982 features with weight zero in the Random Forest regression model, and these features account for around 95.014% of the total number of features (SNPs) (Sup_Table2.). Table 2 showed the top 20 features with higher importance in both the positive side and negative side.

**Table 2.** Top 20 features with higher importance from permutation analysis

|  |  | positive weight |  |  |  | negative weight |  |  |  |
| --- | --- | --- | --- | --- | --- | --- | --- | --- | --- |
| rs# | weight | Std** | P-value | score | rs# | weight | Std** | P-value | score |
| ss715636302 | 0.006481 | 0.000887 | 1 | 0 | ss715584181 | -0.01309 | 0.006558 | 1 | 5.19E-05 |
| ss715632046 | 0.006242 | 0.000954 | 1 | 0 | ss715606461 | -0.01298 | 0.003566 | 1 | 1.08E-05 |
| ss715586408 | 0.005429 | 0.001522 | 1 | 4.26E-05 | ss715611174 | -0.01231 | 0.008024 | 1 | 7.61E-06 |
| ss715629729 | 0.005398 | 0.00163 | 1 | 3.52E-07 | ss715601294 | -0.01182 | 0.002501 | 1 | 1.57E-05 |
| ss715639086 | 0.005334 | 0.001063 | 1 | 0 | ss715621258 | -0.01084 | 0.006296 | 1 | 2.42E-06 |
| ss715600775 | 0.0051 | 0.001249 | 1 | 1.62E-05 | ss715636617 | -0.01053 | 0.00258 | 1 | 0 |
| ss715634776 | 0.004993 | 0.000713 | 1 | 0 | ss715614457 | -0.01017 | 0.004173 | 1 | 5.59E-06 |
| ss715621250 | 0.004987 | 0.002158 | 1 | 2.42E-06 | ss715584279 | -0.00995 | 0.002648 | 1 | 5.13E-05 |
| ss715606657 | 0.004812 | 0.002458 | 0.499356 | 1.06E-05 | ss715616125 | -0.00898 | 0.001576 | 1 | 4.58E-06 |
| ss715610241 | 0.004781 | 0.001113 | 1 | 8.27E-06 | ss715638986 | -0.00891 | 0.007572 | 1 | 0 |
| ss715597582 | 0.004612 | 0.0013 | 1 | 1.97E-05 | ss715619437 | -0.00862 | 0.002983 | 1 | 3.08E-06 |
| ss715611890 | 0.004579 | 7.37E-05 | 1 | 7.16E-06 | ss715610413 | -0.00722 | 0.002548 | 0.263147 | 8.13E-06 |
| ss715632589 | 0.004394 | 0.004083 | 1 | 0 | ss715604675 | -0.00668 | 0.00313 | 1 | 1.23E-05 |
| ss715580947 | 0.004045 | 0.0068 | 1 | 8.9E-05 | ss715605467 | -0.00666 | 0.00415 | 1 | 1.17E-05 |
| ss715596598 | 0.003836 | 0.003571 | 1 | 2.09E-05 | ss715607569 | -0.00632 | 0.006535 | 1 | 9.93E-06 |
| ss715617936 | 0.003701 | 0.002292 | 0.846353 | 3.72E-06 | ss715588364 | -0.00615 | 0.001517 | 1 | 3.63E-05 |
| ss715579007 | 0.003535 | 0.001199 | 1 | 0.00025 | ss715590471 | -0.00558 | 0.001448 | 1 | 3.1E-05 |
| ss715609664 | 0.003281 | 0.001464 | 1 | 8.63E-06 | ss715598227 | -0.00557 | 0.005702 | 1 | 1.89E-05 |
| ss715625697 | 0.003213 | 0.000729 | 1 | 1.09E-06 | ss715632123 | -0.00539 | 0.0025 | 1 | 0 |
| ss715586600 | 0.003188 | 0.000131 | 1 | 4.18E-05 | ss715607399 | -0.0051 | 0.001592 | 1 | 1.01E-05 |

This table shows the top 20 features with higher importance in both the positive side and negative side from permutation analysis.
\*rs# refers to Reference SNP cluster ID
Weight indicates the feature importance weight of SNP by permutation analysis
\*\* std refers to Standard Deviation
Score indicates the score of each feature by Random Forest score analysis
P-value is calculated by the GAPIT software

Besides the permutation feature importance, the feature importance was also computed by feature scores. The computation of feature scores was implemented by the Random Forest algorithm. Our results showed that there are 36077 features in total got a score bigger than 1E-07. Among them, 33121 features (SNPs) got a score bigger than 1E-06, 19735 features (SNPs) got a score bigger than 1E-05, and 1472 features (SNPs) got a score bigger than 0.0001. A total of 6003 features got a score zero, and these features accounts for 12.466% of the total features (SNPs) (Table 2, Sup_Table3).

### 3. Comparison of different methods for feature importance analysis

As mentioned in above, there were 36077 features (SNPs) identified by Random Forest score analysis and 974 features (SNPs) in total had weight value above zero identified by permutation analysis. Among these 974 positive features (SNPs), there were 806 features (SNPs) confirmed by Random Forest score analysis. There were 1124 features (SNPs) in total got negative weight values identified by permutation analysis. Among these 1124 negative features (SNPs), there were 964 features confirmed by Random Forest score analysis. In total, there were 1770 features (SNPs) confirmed by both of Random Forest score analysis ad permutation analysis. Among these 1770 features (SNPs), there were 146 features (SNPs) with P-value < 1 (69 positives and 77 negatives) (Fig. 2, Sup_Table4.).

**Fig. 2.**
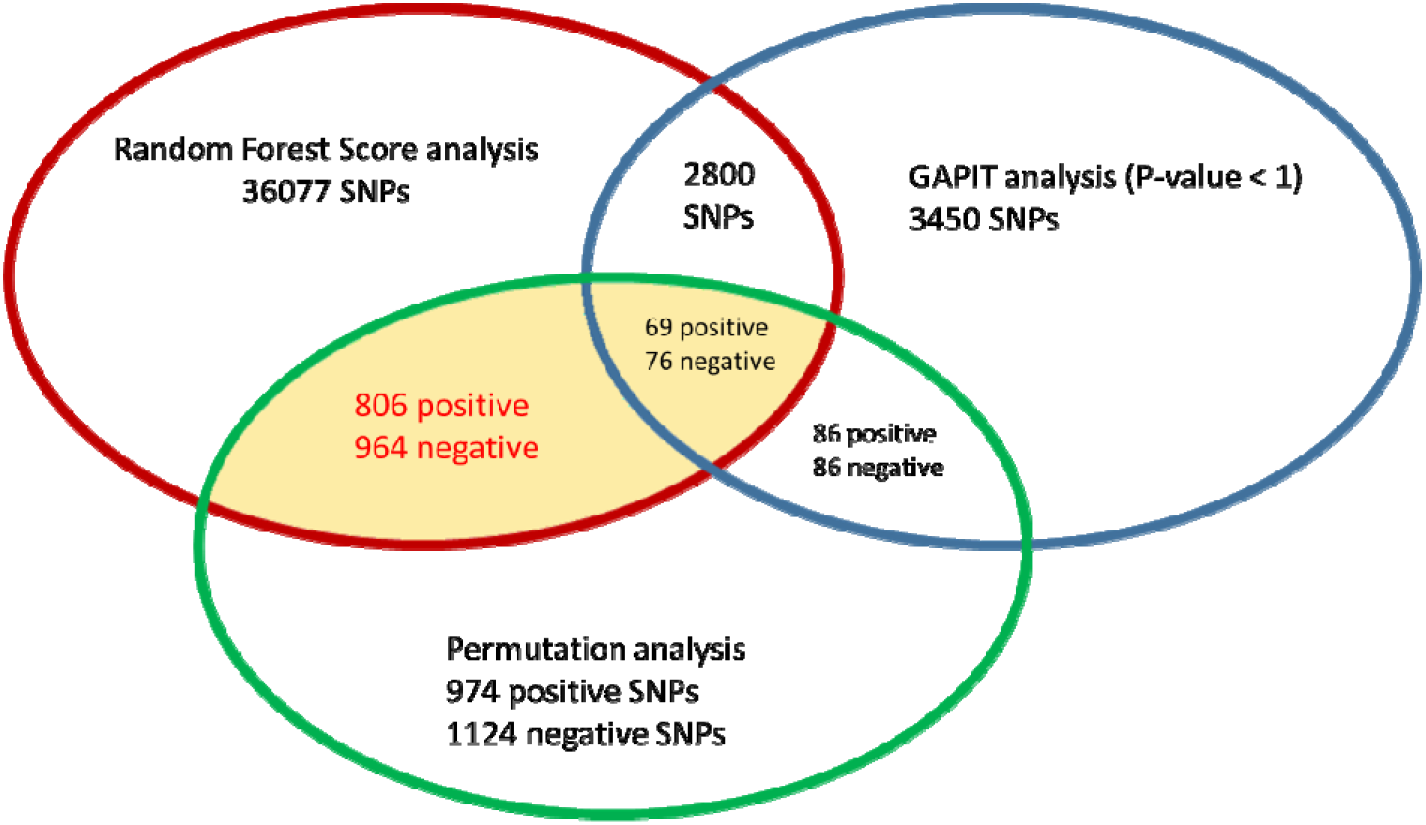
The summary chart of feature importance analysis. This shows a summary of feature importance analysis by the different methods. Blue circle refers to the SNPs with P-value less than 1 identified by GAPIT software; and a total of 3450 SNPs were identified. Green circle refers to the 974 SNPs with positive weight and 1124 SNPs with negative weight, identified by permutation importance analysis. Red circle refers the 36077 SNPs, identified by Random Forest score (score >= 1E-04). The numbers inside the intersection refers to the SNPs, confirmed by both methods. 2800 SNPs were identified by Random Forest score analysis, with P-value less than 1. A total of 1770 SNPs (with 806 SNPs with positive weight and 964 SNPs with negative weight) were confirmed by both of Random Forest score analysis and permutation analysis (highlighted in yellow). 86 SNPs with positive weight and 86 SNPs with negative weight were identified by permutation analysis, with P-value less than 1. The 69 SNPs with positive weight and 76 SNPs with negative weight were confirmed by both of Random Forest score analysis and permutation analysis, with P-value less than 1(highlighted in yellow).

To validate our feature importance analysis results, all 2137 samples characterized with 1170 identified SNPs were applied on the Elastic net regression analysis. Our results showed that the RMSE (root mean square error) was 0.2813 and the R^2^ value was 0.741. Compare to the Elastic net analysis on data subsets from the GAPIT analysis, the accurate level close to the data set those with P-value <1. For SNPs with P-value less than 1 in the GAPIT analysis, the RMSE value was 0.2601 and the R^2^ value was 0.7810, but there were 3451 features (SNPs) applied (Table 2). In other words, our results showed that 1770 features (SNPs) from feature selection could reach the same accuracy as the 3451 features (SNPs) with P-value less than 1.0. The analysis showed that feature importance analysis could help lower the feature size and increase the computation efficiency.

Based on the above analysis, we searched all 1170 SNPs which were confirmed by both of Random Forest score and Permutation analysis in soybean genome. We found that 328 SNPs hit the annotated genes (Sup_Table4). To identify biological processes these 328 genes participate in, we further applied the GO (gene ontology) term enrichment analysis for all of them. Our result showed that the functional group for biological process, cellular component and molecular function were highly enriched by most of these 328 genes (Fig. 3, 4, and 5, Sup_Table5). In biological process, 66 genes (times) were classified into 16 GO term classes and 14 genes had no specific GO term to assign (Fig. 3, Sup_Table5). In cellular component class, 388 genes (times) were classified into 18 GO classes and 14 genes had no specific GO term to assign (Fig. 4, Sup_Table5). In molecular function class, 264 genes (times) were classified into 17 GO classes and 14 genes had no specific GO term to assign (Fig. 5, Sup_Table5). As is common with GO analysis, some genes were classified differently under different GO terms (Sup_Table5).

**Fig. 3.**
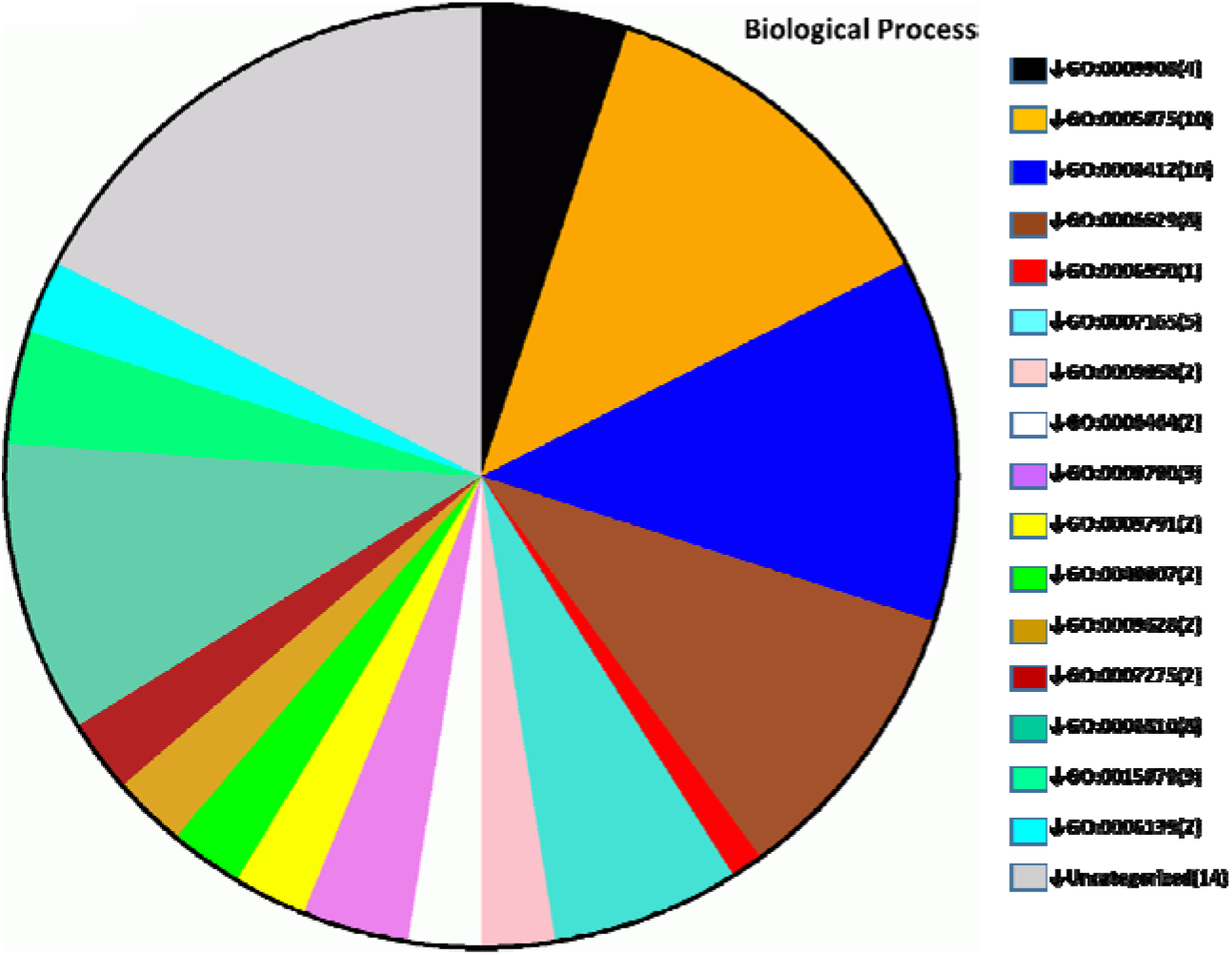
Biological Process Classification. This shows the biological process classification based on GO enrichment analysis. There are 66 genes were classified into 16 GO classes, they are GO:0009908 (Flower Development), GO:0005975(Carbohydrate Metabolic Process), GO:0006412(Translation), GO:0006629 (Lipid Metabolic Process), GO:0006950(Response To Stress), GO:0007165(Signal Transduction), GO:0009058(Biosynthetic Process), GO:0006464 (Protein Modification Process), GO:0009790 (Embryo Development), GO:0009791 (Post-embryonic Development), GO:0040007 (Growth), GO:0009628 (Response To Abiotic Stimulus), GO:0007275 (Multicellular Organismal Development), GO:0006810(Transport), GO:0015979 (Photosynthesis), GO:0006139 (Nucleobase, Nucleoside, Nucleotide And Nucleic Acid Metabolic Process) and there are 14 genes uncategorized. The corresponding gene number is showed in brackets.

**Fig. 4.**
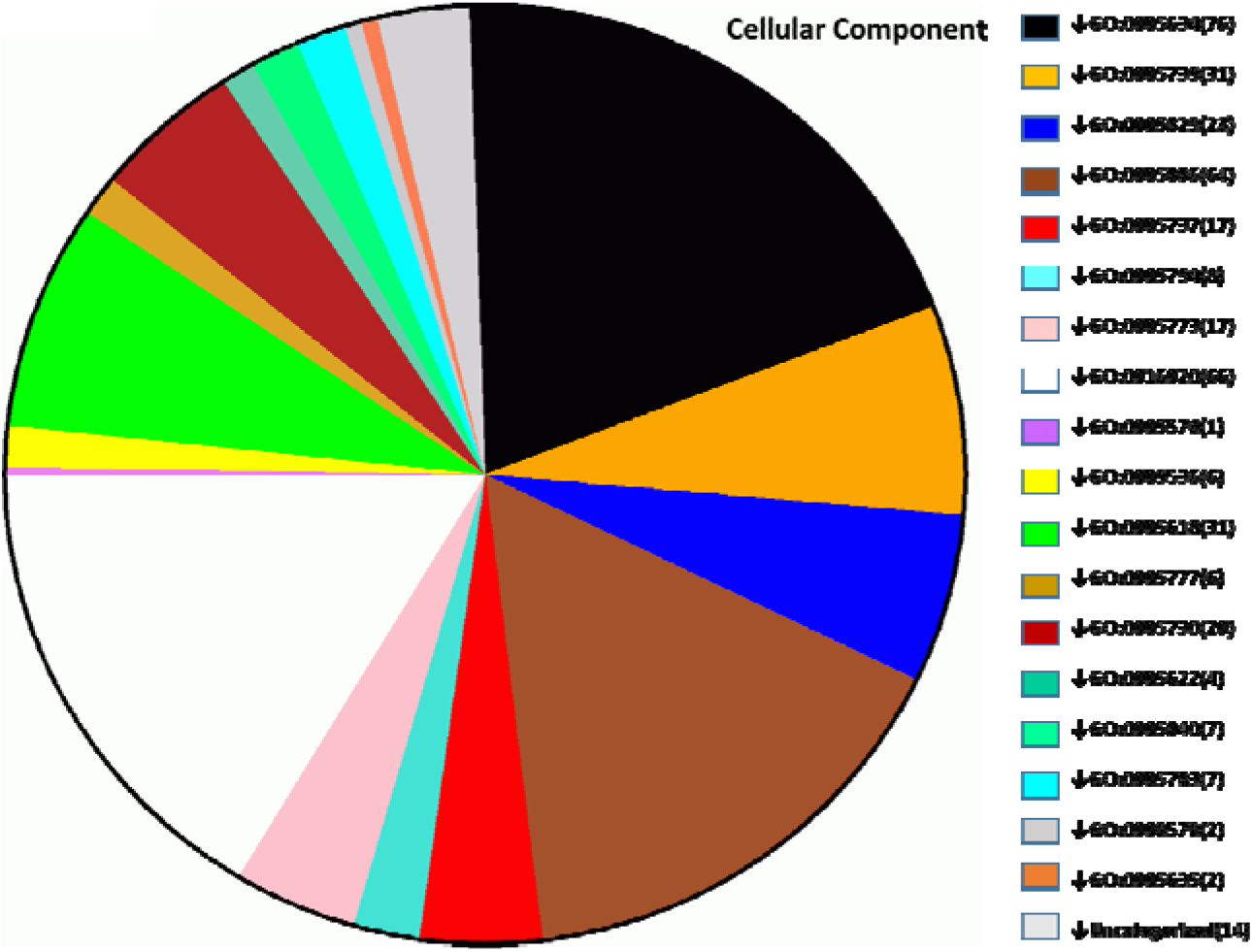
Cellular Component Classification. This shows the cellular component classification based on GO enrichment analysis. There are 388 genes were classified into 18 GO classes, they are GO:0005634(nucleus), GO:0005739(mitochondrion), GO:0005829(cytosol), GO:0005886(plasma membrane), GO:0005737(cytoplasm), GO:0005794(Golgi apparatus), GO:0005773(vacuole), GO:0016020(membrane), GO:0005576(extracellular region), GO:0009536(plastid), GO:0005618(cell wall), GO:0005777(peroxisome), GO:0005730(nucleolus), GO:0005622(intracellular), GO:0005840(ribosome), GO:0005783(endoplasmic reticulum), GO:0009579(thylakoid), GO:0005635(nuclear envelope) and 14 uncategorized. The corresponding gene number is showed in brackets.

**Fig. 5.**
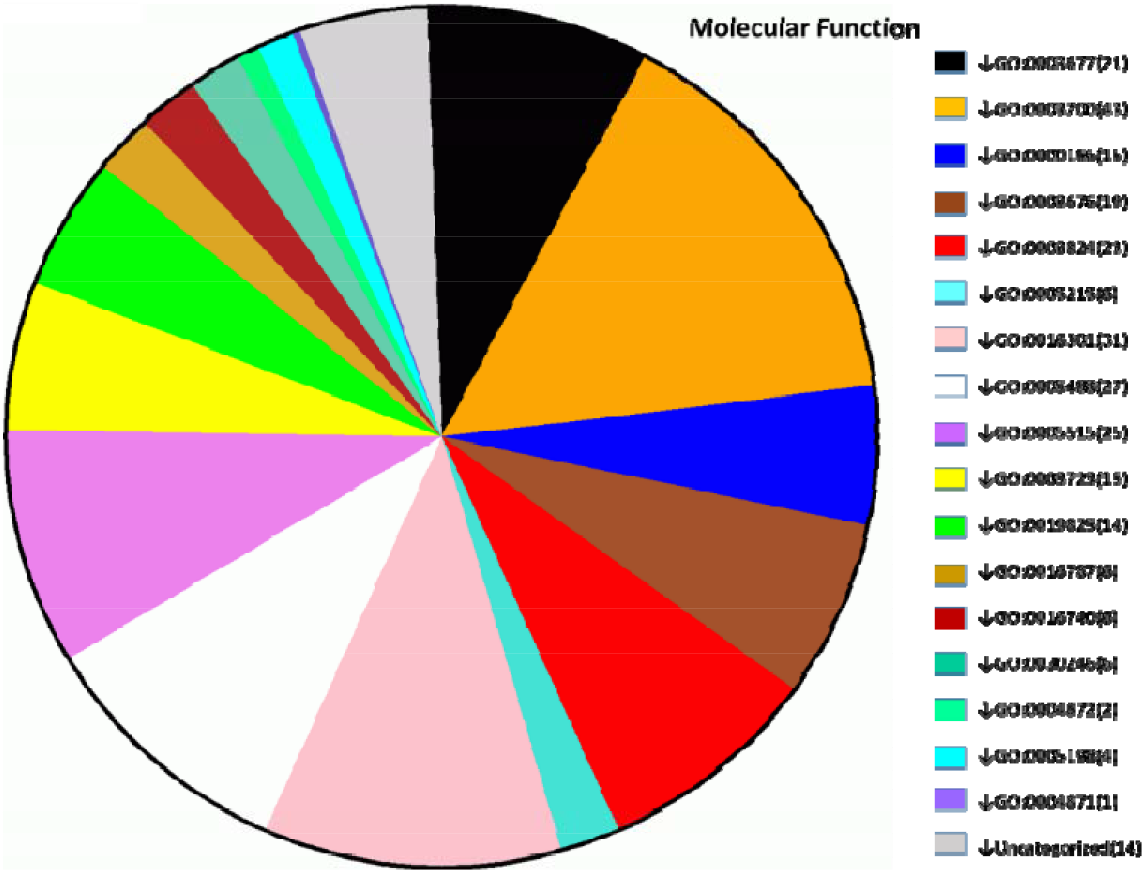
Molecular Function Classification. This shows the molecular function classification based on GO enrichment analysis. There are 264 genes were classified into 17 GO classes, they are GO:0003677(DNA binding), GO:0003700(sequence-specific DNA binding transcription factor activity), GO:0000166(nucleotide binding), GO:0003824(catalytic activity), GO:0005215(transporter activity), GO:0016301(kinase activity), GO:0005488(binding), GO:0005515(protein binding), GO:0003723 (RNA binding), GO:0019825(oxygen binding), GO:0016787(hydrolase activity), GO:0016740(transferase activity), GO:0030246(carbohydrate binding), GO:0004872(receptor activity), GO:0005198(structural molecule activity), GO:0004871 (signal transducer activity) and 14 uncategorized. The corresponding gene number is showed in brackets.

Gene location mapping results showed that all of these 328 genes are scattered on chromosome 1 to chromosome 18. There were no branching related genes located in chromosome 19 and chromosome 20 (Fig. 6). The inquiry term “branching” was searched in Soybase and 35 genes were found (Sup_Table4). To make a comparison, the location of these 35 genes were also marked on Fig. 6. The gene expression information of all 328 genes identified in this research were searched against Soybase for a further analysis (Sup_Table 6).

**Fig. 6.**
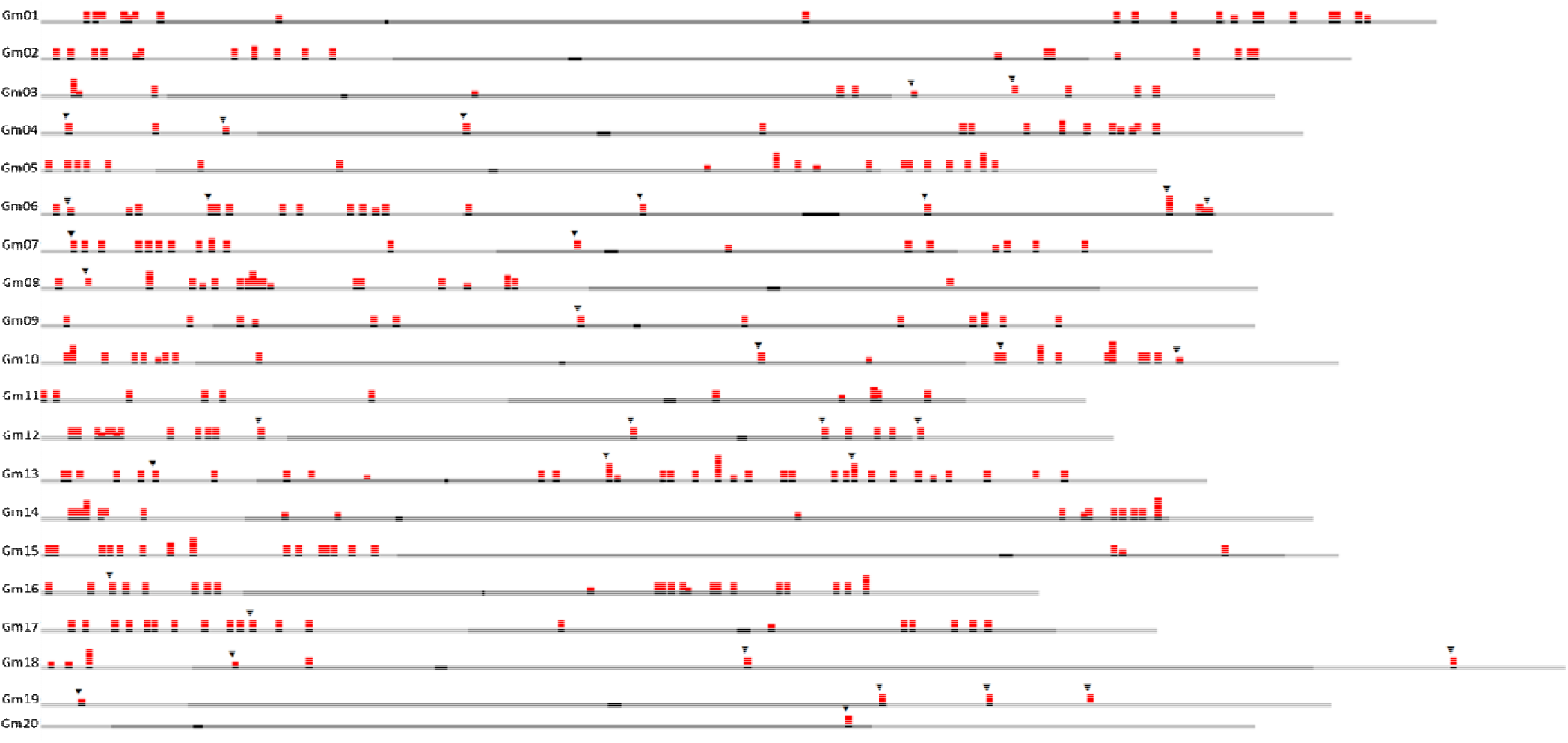
Gene location map. This shows the location of 328 genes, identified by our feature importance analysis. In soybase, there are 35 branching related genes were previously reported; For comparison, the 35 genes are also added to this map (marked by ▼). Color coding is used in the genome viewer to differentiate each query in a multiple FASTA submission. The height of the colored indicators is proportional to the number of BLAST hits in that genomic bin.

## Discussion

### 1. Minor QTLs and genomic selection

Genomic selection is a marker-assisted selection approach to enhance quantitative traits in breeding population, in which whole genome SNPs (single-nucleotide polymorphisms) markers can be used to predict breeding values. Genomic selection has been proved to increase breeding efficiency in both plant and animal breeding, such as dairy cattle, pig, rice and soybean [41]. To get an accurate prediction in genomic selection, we need a better understanding of the population of SNP makers and the contribution of each markers. In the last decade, efforts of global international collaborations have revealed numerous loci that influence traits development in different organism by genotyping and phenotyping very large cohorts of individuals. However, the effects of single alleles explain only a small portion of the heritable variability [42]. Although some traits loci are found, these loci alone do not point to the underlying mechanism responsible for the association, which is due to complex gene interactions in biological activities. To identify genes and pathways responsible for variation in quantitative traits, it is still a central challenge of modern genetics.

Plant breeding is the process of pyramiding favorable alleles. The minor effect QTLs have much more importance in molecular breeding and commercial breeding since the enrichment of minor alleles can enhance the control accuracy of phenotype performance [43]. In this research, we applied a new framework which combined the GWAS analysis and different feature selection methods to explore minor QTLs/alleles and their importance in soybean branching. Compare to the P-value method in GWAS analysis, the feature importance analysis we used in this research explored 36077 features in total with a score higher than 1E-07, which is about ten times as the number of the features identified in GWAS analysis with P-value less than 1.0. Based on the Permutation feature importance analysis, we explored 974 features with positive effects on soybean branching development and 1124 features with negative effects on soybean branching development (Table 2 and Sup_Table2). Either in linkage mapping or in association mapping, it is difficult to find the QTLs which have negative contribution to a trait, even we all know there are negative QTLs/alleles involved in all biological activities. From our analysis and testing results, the new framework we used in this research is superior to the traditional P-value based methods in molecular genetics analysis. Actually, in GWAS analysis, there is only one SNP (ss715607451) above the threshold, unfortunately, this SNP does not hit on any gene. And the BLAST results of the 18 SNPs with P-value less than 0.005 in Soybase shows five annotated genes (Table 1), but none of them is reported as branching related. All of these information are very important to genomic selection and could lead to an accurate prediction in genomic selection a further study in future.

### 2. Feature importance analysis and its applications

In this research, we applied three kind of feature importance analysis, permuted feature importance, feature importance scoring and P-value analysis through GAPIT. We employed the Random Forest regression algorithm in permuted feature importance and feature importance scoring analysis. It is reported that the feature importance based methods are applicable if we are going to use a tree-based model for making predictions [44]. Random Forest is one of the most effective machine learning models for predictive analytics capable of performing both regression and classification tasks and able to capture non-linear interaction between the features and the target [45]. In random Forest, features that tend to split nodes closer to the root of a tree will result in a larger importance value. Node splits based on this feature on average result in a large decrease of node impurity. Permutation feature importance is a model-agnostic approach and is calculated after a model has been fitted. The values of a feature of interest and reevaluate model performance is permutated after evaluating the performance of model. The observed mean decrease in performance indicates feature importance. The performance decrease can be compared on the test set as well as the training set. Only the latter will tell us something about generalizable feature importance.

As we mentioned above, one of the biggest problems facing GWAS analysis is difficult to detect quantitative traits which controlled by multiple genes, Association mapping and bi-parent mapping good for major QTLs but not minor QTLs, Minor QTLs are important for quantitative traits but hard to be detected by traditional genetic research, Machine learning methods open a door for minor QTLs mining, special for non-model organisms with less research basis. Our results showed that the new framework displays much powerful ability in minor QTLs mining than conventional analysis methods. We can expect many discoveries will be made through applications of different machine learning methods to genomics data, particularly in genomic selection research.

## Conclusions

Accurate prediction of genomic breeding values is a central challenge to contemporary plant and animal breeders. Minor QTLs play very important roles in this procedure, but we know little about the minor QTLs in most traits’ development. To understand how many genes and which genes involved in the trait’s development is the prerequisites of breeding prediction. In this research, we combined the GWAS analysis and feature selection with machine learning methods, and explored the new framework in minor QTLs mining. The framework provides a way for finding minor QTLs and better estimates of the QTL effects supportable by the data. Unlike QTL mapping through linkage mapping, this framework does not require a genetic map. It is therefore applicable to any species or population. This research on minor QTLs miming will contribute to trait’s development and gene pathway analysis in further studies.

## Abbreviations

SNPs: single-nucleotide polymorphisms
GWAS: genome-wide association study
CDS: coding region sequence
UTR: untranslated region
GO: gene ontology
BLAST: Basic Local Alignment Search Tool

## Declarations

## Acknowledgments

We thank our campus research colleagues for their helpful suggestions, insightful comments and discussions on this research.

## Availability of data and materials

The original dataset is publically available. And our intermediate analysis results and code used for analysis with this study are available from the corresponding author upon request.

## Authors’ contributions

WZ participated in the statistical analyses, data processing and writing the manuscript. XH participated in conceiving the presented idea, development of the software, discussions of the results, and drafted the manuscript. EB, JS, JC, JQ and KW collaborated with statistical analyses, data processing, interpretation, data analysis support, and writing of the manuscript. All the authors approved the manuscript.

## Ethics approval and consent to participate

NA

## Consent for publication

NA

Seven Additional Files:

Sup_Table1. GAPIT.MLM.Branching.GWAS.Results.csv

Sup_Table2. RF_Perm_importance.xlsx

Sup_Table3. RF_feature_score.xlsx

Sup_Table4. gene Blast result.xls

Sup_Table5. gene ontology analysis.xlsx

Sup_Table6. gene_expression information.csv

Sup_Table7. RS_HeaderT.csv

